# A lactate-HIF1α-VDR positive feedback loop drives protective SPP1⁺ macrophage differentiation to inhibit schistosomiasis-induced liver fibrosis

**DOI:** 10.64898/2026.09.15.751751

**Authors:** Zhou Xing, Pingping Yang, Huiyu Xia, Rui Yin, Bin Le, Fangbin Zhou, Xiaoying Guo, Xiaobin Fan, Xing He

**Affiliations:** Department of Tropical Diseases, Shanghai Key Laboratory of Medical Bioprotection, Key Laboratory of Biological Defense, Ministry of Education, Naval Medical University, Shanghai 200433, China; Department of Neurology, No. 902 Hospital of People’s Liberation Army Joint Logistics Support Force, Bengbu 233000, China; Department of Environmental Health, School of Public Health, China Medical University, Shenyang 110122, Liaoning Province, China

**Author notes:** These authors contributed equally to this work (Zhou Xing, Pingping Yang, Huiyu Xia). Corresponding author. (X.G.); (X. F.); (X.H.).

**Keywords:** schistosomiasis, hepatic fibrosis, macrophage, vitamin D receptor, SPP1

## Abstract

Hepatic fibrosis remains a major cause of morbidity and mortality in patients with schistosomiasis without effective therapy. Here we show that vitamin D receptor (VDR) signaling in macrophages is essential for limiting fibrosis, acting through a previously unrecognized metabolic-epigenetic circuit. Hepatic macrophages exhibit the highest VDR expression among liver-resident cells, and pharmacological VDR activation with paricalcitol selectively expands a protective SPP1⁺ macrophage subset derived from circulating monocytes. Myeloid-specific VDR knockout exacerbates fibrosis and abrogates paricalcitol’s hepatoprotective effects, whereas SPP1⁺ macrophage depletion worsens disease. Mechanistically, VDR activation synergizes with hypoxia and lactate to drive SPP1 expression via glycolytic reprogramming. Moreover, lactate and HIF1α cooperatively induce VDR transcription by binding to the *Vdr* promoter and enhancing histone lactylation, forming a positive feedback loop that amplifies the antifibrotic response. Our findings establish the lactate-HIF1α-VDR-SPP1 axis as an endogenous defense mechanism and identify macrophage VDR as a promising therapeutic target for fibrotic liver diseases.

## INTRODUCTION

Schistosomiasis is a neglected tropical disease caused by infection with blood flukes of the genus Schistosoma, affecting over 250 million people worldwide, predominantly in sub-Saharan Africa (*1*). The major morbidity and mortality of chronic intestinal schistosomiasis result from hepatic fibrosis induced by parasite eggs trapped in the liver, which can progress to portal hypertension, variceal bleeding, and ultimately liver failure (*2*). Despite decades of effort, no effective antifibrotic therapy is currently approved for clinical use, highlighting an urgent need to understand the underlying mechanisms and identify novel therapeutic targets. Hepatic fibrosis in schistosomiasis is fundamentally an immunopathological disease, driven by the host immune response to parasite eggs trapped in the liver (*3*,*4*). Upon egg deposition, a complex inflammatory cascade is initiated, leading to the recruitment and activation of various immune cells. Among these, macrophages play a central and versatile role in the pathogenesis of liver fibrosis. Depending on the signals from the local microenvironment, macrophages can polarize into distinct phenotypic subsets that either promote or resolve fibrosis (*5*). Notably, macrophages have been shown to exert critical functions at all stages of tissue repair and fibrosis, and they are increasingly recognized as a promising cellular target for developing novel antifibrotic therapies (*6*).

The vitamin D receptor (VDR) is a nuclear hormone receptor best known for its role in calcium and bone metabolism (*7*). Recent studies have uncovered its antifibrotic potential. Ding *et al*. demonstrated that VDR ligands inhibit the activation of hepatic stellate cell (HSC), the principal effector cell of hepatic fibrosis, by antagonizing SMAD3 occupancy on profibrotic gene promoters (*8*). In the context of schistosomiasis, our previous work showed that VDR expression in HSCs is negatively regulated by an IFN-γ/IRF2/miR-351 axis, and that activation of VDR signaling attenuates egg-induced hepatic fibrosis (*9*). These findings established VDR as a critical endogenous brake on HSC-driven fibrogenesis. In addition, VDR has also emerged as an important immunomodulator, as it is expressed in various immune cells including monocytes, macrophages, and T lymphocytes, and its activation regulates cytokine production, antigen presentation, and immune cell differentiation (*10-12*).

While the protective role of VDR in HSC is well recognized, our preliminary data revealed that among liver-resident cells, VDR expression is highest in macrophages, prompting us to investigate whether VDR modulates hepatic fibrosis by regulating the hepatic immune microenvironment beyond its cell-autonomous effect on HSCs. Using single-cell RNA sequencing, we observed that pharmacological activation of VDR with paricalcitol markedly ameliorated liver fibrosis and, intriguingly, increased the abundance of SPP1^+^ macrophages in the liver. Myeloid-specific VDR knockout exacerbated fibrosis and reduced SPP1^+^ macrophage accumulation, whereas SPP1⁺ macrophage depletion worsened disease. Mechanistically, VDR activation synergized with hypoxia, HIF1α, and lactate to promote SPP1 expression through inducing glycolytic reprogramming in macrophages. In turn, HIF1α and lactate acted in concert to promote VDR transcription through binding to the *Vdr* promoter and enhancing histone lactylation. Collectively, these findings reveal a previously unrecognized mechanism by which VDR signaling in macrophages drives their differentiation into an SPP1^+^ phenotype, thereby limiting pathological fibrogenesis.

## RESULTS

### VDR activation ameliorates schistosomiasis-induced hepatic fibrosis and remodels the liver immune microenvironment

To investigate the cellular basis of VDR-mediated antifibrotic effects in schistosomiasis, we first examined VDR expression in major liver-resident cell populations. Quantitative PCR (qPCR) analysis revealed that among hepatocytes, macrophages, HSCs, liver endothelial sinusoidal cells (LESCs), and CD45^+^ immune cells, VDR mRNA levels were highest in macrophages and CD45^+^ cells (Supplementary Fig. 1A). Furthermore, treatment of isolated hepatic macrophages with the VDR agonist paricalcitol significantly induced the expression of CYP24A1, a well-established VDR target gene, indicating that hepatic macrophages are functionally responsive to VDR activation (Supplementary Fig. 1B).

We next evaluated the therapeutic effect of paricalcitol in a mouse model of *S. japonicum*-induced hepatic fibrosis. C57BL/6 male mice were percutaneously infected with cercariae, and then treated with paricalcitol or vehicle three times per week. At day 42 post-infection, when robust granuloma formation and fibrosis had developed, paricalcitol-treated mice exhibited a marked reduction in hepatic hydroxyproline content, a biochemical indicator of collagen deposition (Supplementary Fig. 2A). Histological analysis confirmed that paricalcitol treatment significantly decreased both the size of egg-induced granulomas (Supplementary Fig. 2B, D) and the liver fibrosis score (Supplementary Fig. 2C, D). Consistent with these observations, mRNA levels of fibrotic markers including Col1a1, Timp1, and Acta2 were substantially downregulated in the paricalcitol group (Supplementary Fig. 2E). Moreover, the markers of hepatocyte injury, including serum alanine aminotransferase (ALT) and aspartate aminotransferase (AST) levels, were significantly reduced upon paricalcitol treatment (Supplementary Fig. 2F). Importantly, no significant differences were observed in worm pair counts or hepatic egg burden between vehicle- and paricalcitol-treated mice (Supplementary Fig. 2G), indicating that the antifibrotic effect of paricalcitol was not mediated by altering parasite burden. Collectively, these data demonstrate that pharmacological activation of VDR signaling effectively attenuates schistosomiasis-induced hepatic pathology.

To dissect the impact of VDR activation on the liver immune microenvironment, we performed single-cell RNA sequencing (scRNA-seq) of hepatic CD45^+^ immune cells isolated from uninfected mice, infected vehicle-treated mice, and infected paricalcitol-treated mice at day 42 post-infection (Fig. 1A). After quality control, a total of 21,073 cells were obtained. Unsupervised clustering and uniform manifold approximation and projection (UMAP) visualization identified eight major immune cell types based on canonical marker genes: B cells, neutrophils, monocytes/macrophages, Kupffer cells, T cells, NK cells, dendritic cells, and basophils (Fig. 1B and 1C). Comparative analysis revealed striking alterations in the immune landscape upon infection and VDR activation. Specifically, infection increased the proportions of neutrophils and monocytes/macrophages, whereas paricalcitol treatment reversed these changes. Conversely, the reduced proportions of B cells, T cells, and NK cells following infection were partially restored by paricalcitol (Fig. 1D-F). We next calculated a VD response score for each cell based on the expression of VDR pathway related genes. Interestingly, the overall VD response score in hepatic CD45^+^ cells was elevated after infection and further increased upon paricalcitol treatment (Fig. 1G). Among the eight immune subsets, monocytes/macrophages exhibited the highest VD response score (Fig. 1H and 1I). These results strongly suggest that monocytes/macrophages are the primary cellular targets of paricalcitol within the liver immune microenvironment, prompting us to focus on this population for subsequent mechanistic investigations.

**Figure 1.**
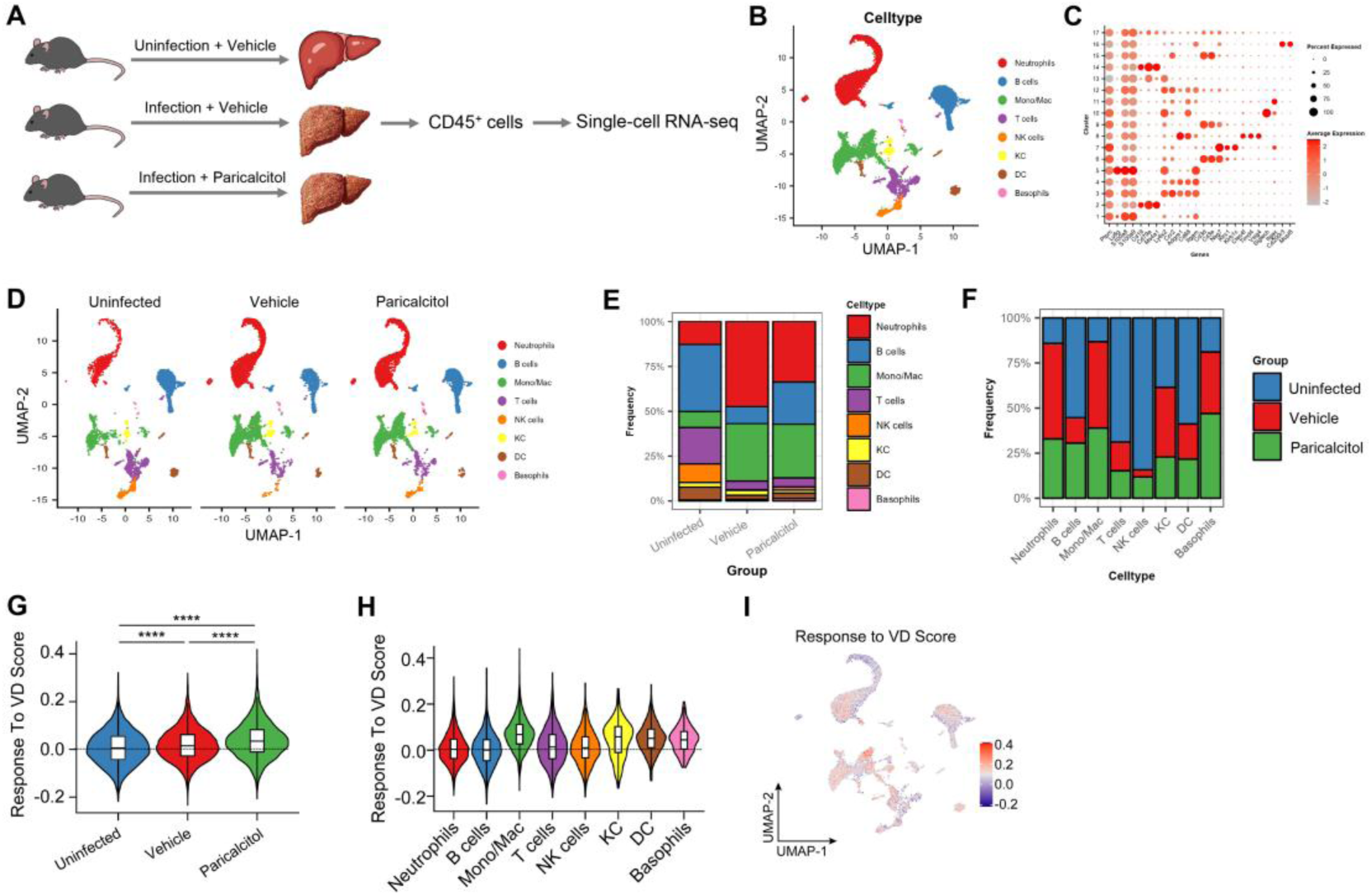
Single-cell transcriptomic analysis reveals that monocytes/macrophages are the primary targets of paricalcitol in the liver during schistosomiasis. (**A**) Experimental workflow: Uninfected mice, infected vehicle-treated mice, and infected paricalcitol-treated mice (n = 5 per group) were sacrificed at day 42 post-infection. Hepatic CD45⁺ cells were isolated from each group, pooled within the same group, and subjected to single-cell RNA sequencing. (**B**) UMAP plot of 21,073 CD45^+^ immune cells colored by cell type annotation based on canonical marker genes. (**C**) Heatmap showing expression of representative marker genes for each identified cell type. (**D**) UMAP plots of cells from uninfected, infected vehicle-treated, and infected paricalcitol-treated groups, colored by cell type. (**E**) Stacked bar plot showing the frequency of hepatic immune cell subsets in the three groups. (**F**) Stacked bar plot showing the group-specific frequency distribution of distinct immune cell populations. (**G**) Violin plot showing the VD response score in total CD45^+^ cells across the three groups. (**H**) Violin plots showing VD response scores for each cell type. (**I**) UMAP plot showing VD response scores across individual cells. Data from panels G and H are presented as violin plots, with embedded boxplots showing the median (center line) and interquartile range (box boundaries). The width of each violin represents the kernel density estimation of the data distribution. Statistical analysis was performed using the Kruskal-Wallis, followed by Dunn’s post-hoc test with Bonferroni correction. \*\*\*\**p*<0.0001.

### VDR activation selectively expands a distinct SPP1^+^ macrophage subset that exhibits the highest vitamin D response score

To further dissect the heterogeneity of monocytes/macrophages and identify the specific subset(s) targeted by paricalcitol, we performed subclustering analysis of all monocyte/macrophage cells from the scRNA-seq dataset. Unsupervised clustering based on differential gene expression revealed eight distinct subclusters: Ly6Chi monocytes, Mertk^+^ macrophages, SPP1^+^ macrophages, Ly6Clo monocytes 1, Ltk macrophages, proliferating monocytes, Ly6Clo monocytes 2, and unknown monocytes (Fig. 2A and 2B). Except for the unknown monocyte cluster, all other subsets expanded upon schistosome infection (Fig. 2C-E). Strikingly, while paricalcitol treatment reduced the proportions of most monocyte/macrophage subsets compared to the vehicle-treated infected group, the SPP 1^+^ macrophage subset displayed a unique behavior: its numbers further increased upon paricalcitol treatment (Fig. 2C-E). Notably, SPP 1^+^ macrophages were virtually absent in uninfected livers and emerged specifically upon infection, confirming their infection-associated identity. We next calculated the VD response score for each monocyte/macrophage subset. The overall VD response score in total monocytes/macrophages was elevated after infection and further increased by paricalcitol treatment (Fig. 2F). Among the eight subsets, SPP1^+^ macrophages exhibited the highest VD response score (Fig. 2G and 2H), suggesting that this subset is the primary responder to VDR activation.

**Figure 2.**
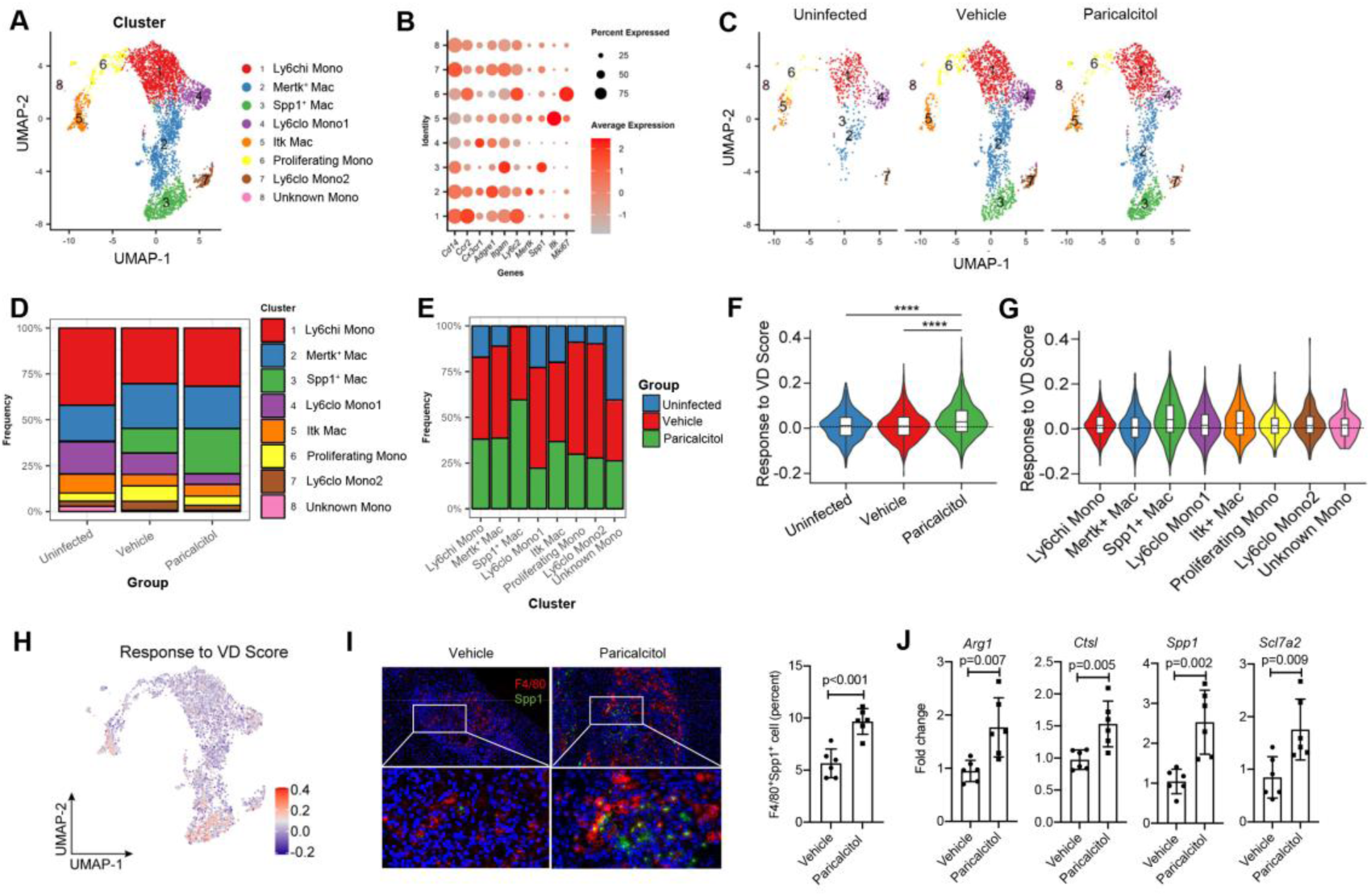
SPP1^+^ macrophage is the predominant paricalcitol-responsive subset within the hepatic monocyte/macrophage pool. (**A**) UMAP plot of monocyte/macrophage subclusters colored by subcluster annotation based on marker genes. (**B**) Heatmap showing expression of representative marker genes for each subcluster. (**C**) UMAP plots of monocyte/macrophage subclusters from uninfected, infected vehicle-treated, and infected paricalcitol-treated groups. (**D**) Stacked bar plot showing the frequency of hepatic monocyte/macrophage subclusters in the three groups. (**E**) Stacked bar plot showing the group-specific frequency distribution of distinct hepatic monocyte/macrophage subclusters. (**F**) Violin plot showing the VD response score in total monocytes/macrophages across the three groups. (**G**) Violin plots showing VD response scores for each monocyte/macrophage subcluster. (**H**) UMAP plot showing VD response scores across individual monocyte/macrophage cells. (**I**) Representative multiplex immunohistochemistry images of liver sections stained for F4/80 (red), SPP1 (green), and DAPI (blue). Scale bar, 50 μm. (**J**) Relative mRNA expression of SPP1^+^ macrophage signature genes (Arg1, Slc7a2, Spp1, and Ctsl) in F4/80^+^ magnetic bead-purified hepatic macrophages from vehicle- and paricalcitol-treated infected mice, determined by qPCR. Data from panels F and G are presented as violin plots, with embedded boxplots showing the median (center line) and interquartile range (box boundaries). The width of each violin represents the kernel density estimation of the data distribution. Statistical analysis was performed using the Kruskal-Wallis, followed by Dunn’s post-hoc test with Bonferroni correction. \*\*\*\**p*<0.0001. Data from panels J and K are presented as mean ± SD. Statistical analysis was performed using two-tailed Student’s *t*-test. These experiments were performed in triplicate.

To validate these findings, we performed multiplex immunohistochemistry for F4/80 (a pan-macrophage marker) and SPP1. In line with the scRNA-seq data, livers from paricalcitol-treated infected mice showed a marked increase in F4/80^+^SPP1^+^ double-positive cells compared to vehicle-treated infected mice, and these double-positive cells were predominantly localized within egg granulomas (Fig. 2I). Furthermore, we isolated hepatic macrophages using F4/80 magnetic beads and quantified the expression of four signature genes of SPP1^+^ macrophages (Arg1, Slc7a2, SPP1, and Ctsl; Supplementary Fig. 3) by qPCR. Consistent with the single-cell results, paricalcitol treatment significantly upregulated all four signature genes in bulk hepatic macrophages (Fig. 2J). Collectively, these data identify SPP1^+^ macrophages as an infection-induced, paricalcitol-responsive subset, positioning this macrophage subset as the primary cellular mediator of VDR’s antifibrotic effect in schistosomiasis.

### Myeloid-specific VDR knockout aggravates liver fibrosis and abrogates the SPP1^+^ macrophage differentiation

To definitively establish the cell-autonomous role of macrophage VDR in schistosomiasis-induced hepatic pathology, we generated myeloid-specific VDR knockout mice (VDR^lyz2-Cre^) by crossing VDR^fl/fl^ mice with Lyz2-Cre transgenic mice (Supplementary Fig. 4A). Genotyping confirmed successful recombination, and western blot analysis of isolated hepatic macrophages verified efficient VDR deletion specifically in macrophages (Supplementary Fig. 4B and 4C).

We next subjected VDR^fl/fl^ (control) and VDR^lyz2-Cre^ (myeloid-specific VDR knockout) mice to *S. japonicum* infection, followed by treatment with vehicle or paricalcitol. At day 42 post-infection, livers, serum, and hepatic macrophages were collected for analysis. In VDR^fl/fl^ control mice, paricalcitol treatment significantly reduced hepatic hydroxyproline content (Fig. 3A), granuloma size (Fig. 3B and 3D), liver fibrosis score (Fig. 3C and 3D), mRNA levels of fibrotic markers (Fig. 3E), and serum ALT and AST levels (Fig. 3F), while increasing both the proportion of SPP1^+^ macrophages (Fig. 3H) and SPP1 expression in macrophages (Fig. 3I), consistent with our previous observations. Strikingly, in VDR^lyz2-Cre^ mice, macrophage VDR deletion alone led to exacerbated liver pathology, as evidenced by increased hydroxyproline content, granuloma size, fibrosis score, fibrotic gene expression, and ALT or AST levels, compared to infected VDR^fl/fl^ vehicle-treated mice (Fig. 3A-F). Moreover, VDR deletion reduced both the proportion of SPP1^+^ macrophages and SPP1 expression (Fig. 3H and 3I). Importantly, while paricalcitol treatment in VDR^lyz2-Cre^ mice still reduced hydroxyproline content, fibrosis score, and fibrotic gene expression (Fig. 3A, C-E), its beneficial effects on granuloma size, ALT or AST levels, and SPP1^+^ macrophage differentiation were completely abolished (Fig. 3B, D, F, H, I). Of note, no significant differences in worm pair counts or hepatic egg burden were observed among any of the groups (Fig. 3G), indicating that the observed phenotypes were not secondary to altered parasite burden.

**Figure 3.**
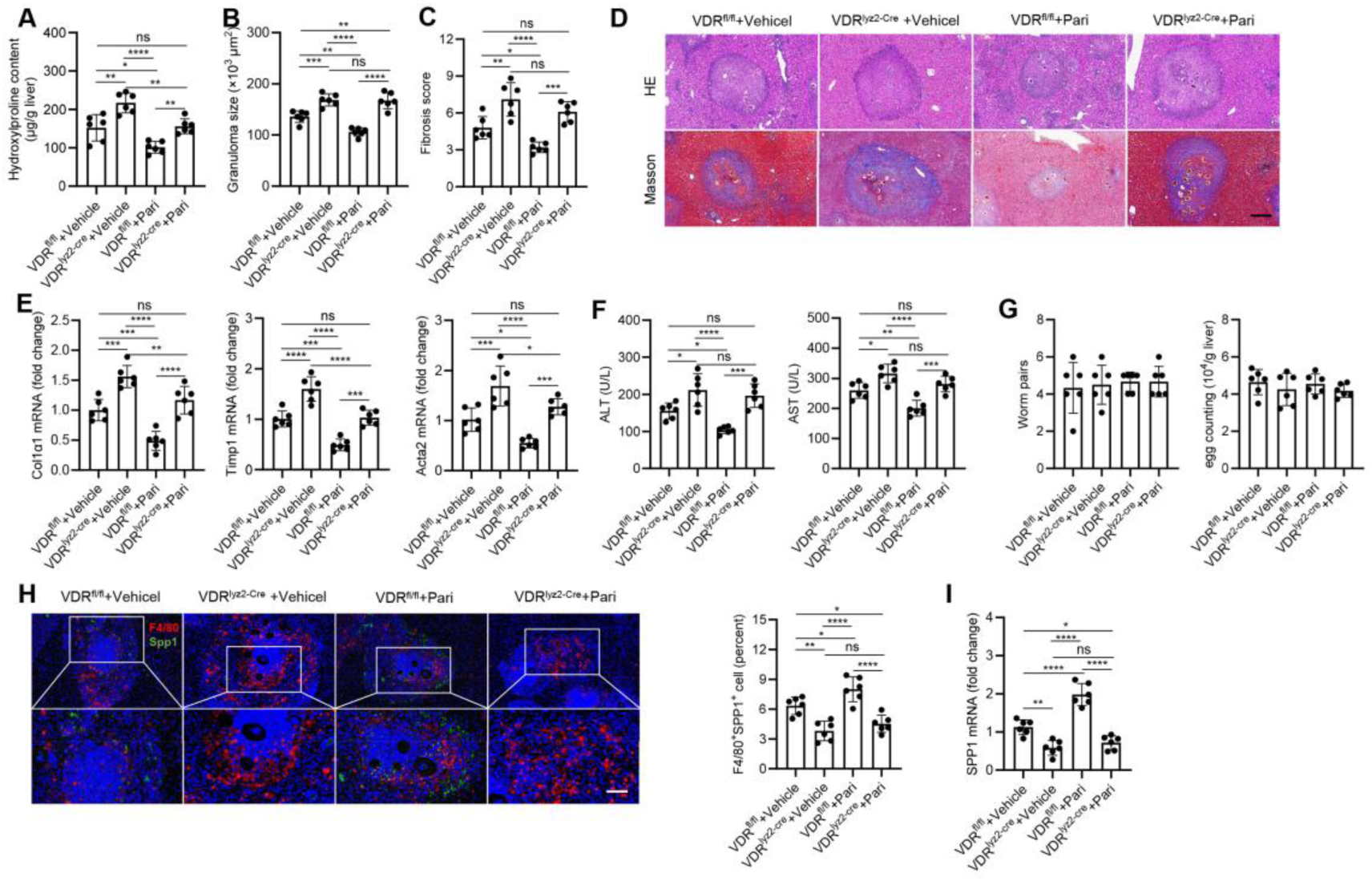
Myeloid-specific VDR knockout exacerbates schistosomiasis-induced liver pathology and abrogates paricalcitol-induced SPP1^+^ macrophage differentiation. VDR^fl/fl^ (control) and VDR^lyz2-Cre^ (myeloid-specific VDR knockout) mice were infected with *S. japonicum* cercariae and treated with vehicle or paricalcitol three times per week. Livers, serum, and hepatic macrophages were collected at day 42 post-infection. (**A**) Hepatic hydroxyproline content. (**B**) Granuloma size measured from H&E-stained liver sections. (**C**) Fibrosis score assessed by Masson’s trichrome staining. (**D**) Representative images of Masson’s trichrome staining (upper) and H&E staining (lower) of liver sections. Scale bar, 200 μm. (**E**) Relative mRNA expression of fibrotic markers Col1a1, Timp1, and Acta2 in liver tissues determined by qPCR. (**F**) Serum ALT and AST levels. (**G**) Adult worm pairs and hepatic egg burden. (**H**) Representative multiplex immunohistochemistry images of liver sections stained for F4/80 (red), SPP1 (green), and DAPI (blue). Scale bar, 50 μm. **(I**) Relative mRNA expression of SPP1 in F4/80^+^ magnetic bead-purified hepatic macrophages determined by qPCR. Data are presented as mean ± SD. Statistical analysis was performed using ANOVA followed by Tukey’s post hoc test. The experiments were performed in triplicate.

Collectively, these results demonstrate that macrophage VDR is essential for the paricalcitol-mediated protection against granulomatous inflammation, hepatocyte injury, and SPP1^+^ macrophage differentiation, but its role in suppressing fibrotic gene expression may involve additional paricalcitol-responsive cell types, such as HSCs. These data firmly establish macrophage VDR as a critical regulator of the protective SPP1^+^ macrophage phenotype during schistosomiasis.

### Depletion of SPP1^+^ macrophages exacerbates schistosomiasis-induced hepatic fibrosis

To directly determine the functional contribution of SPP1^+^ macrophages to liver pathology, we generated SPP1^+^ macrophage reporter/depletion mice by crossing SPP1^LSL-DTR-EGFP^ mice with Lyz2-Cre mice (Supplementary Fig. 5A). In this system, Cre-mediated excision of the loxp-flanked stop cassette enables expression of DTR (diphtheria toxin receptor) and EGFP specifically in SPP1^+^ macrophages, allowing both visualization and DT-mediated depletion. Genotyping confirmed proper recombination (Supplementary Fig. 5B). Flow cytometry analysis showed that DT treatment effectively reduced EGFP^+^ cells in SPP1^LSL-DTR-EGFP/+^Lyz2-Cre^+/-^ mice (Supplementary Fig. 5C).

We then infected SPP1^LSL-DTR-EGFP/+^Lyz2-Cre^+/-^ (SPP1-DTR) and control SPP1^LSL-DTR-EGFP/+^ (SPP1-LSL) mice with *S. japonicum* cercariae. Starting at day 21 post-infection, mice received either DT or PBS. At day 42, livers and serum were collected. Depletion of SPP1^+^ macrophages significantly aggravated liver fibrosis, as evidenced by increased hepatic hydroxyproline content (Fig. 4A), elevated fibrosis score (Fig. 4B and 4C), and enhanced expression of fibrotic marker genes Col1a1, Timp1, and Acta2 (Fig. 4E). Granuloma size was also markedly enlarged in DT-treated SPP1-DTR mice (Fig. 4B and 4D). Serum ALT and AST levels, indicators of hepatocyte injury, were elevated upon SPP1^+^ macrophage depletion (Fig. 4F). No significant differences in parasite burden were observed (Fig. 4G). Furthermore, DT had no effect on liver pathology and parasite burden in control mice (Fig. 4A-G). Collectively, these data demonstrate that SPP1^+^ macrophages play a protective, antifibrotic role during schistosomiasis-induced liver pathology.

**Figure 4.**
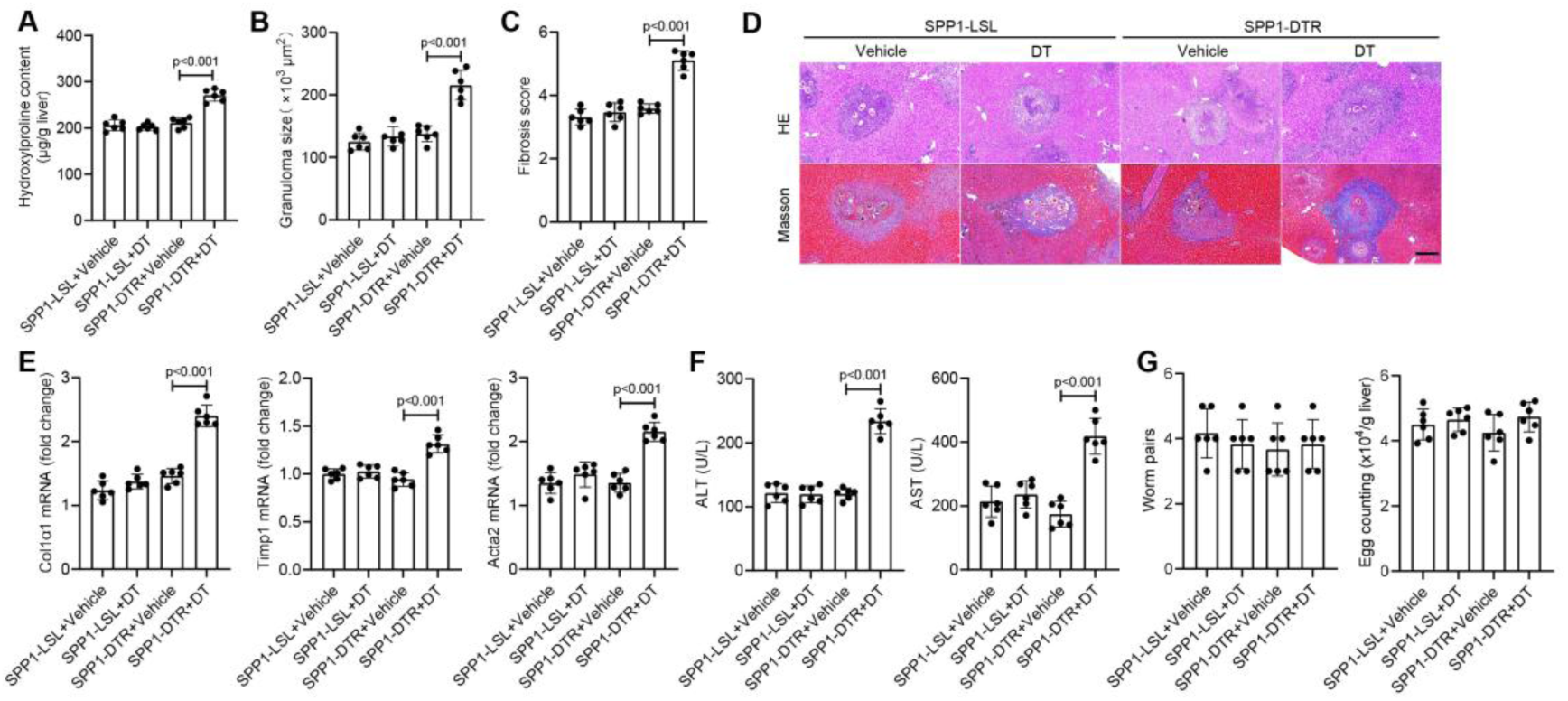
Depletion of SPP1^+^ macrophages aggravates schistosomiasis-induced hepatic fibrosis. SPP1^LSL-DTR-EGFP/+^Lyz2-Cre^+/-^ (SPP1-DTR) and SPP1^LSL-DTR-EGFP/+^Lyz2-Cre^-/-^ (control) mice were infected with *S. japonicum* cercariae. DT or PBS was administered intraperitoneally starting at day 21 post-infection. Livers and serum were collected at day 42. (**A**) Hepatic hydroxyproline content. (**B**) Representative images of Masson’s trichrome staining (upper) and H&E staining (lower) of liver sections. Scale bar, 200 μm. (**C**) Fibrosis score assessed by Masson’s trichrome staining. (**D**) Granuloma size measured from H&E-stained liver sections. (**E**) Relative mRNA expression of fibrotic markers Col1a1, Timp1, and Acta2 in liver tissues determined by qPCR. (**F**) Serum ALT and AST levels. (**G**) Adult worm pairs and hepatic egg burden. Data are presented as mean ± SD. Statistical analysis was performed using ANOVA followed by Tukey’s post hoc test. The experiments were performed in triplicate.

### SPP1^+^ macrophages originate from circulating monocytes and require hypoxia/glycolysis for terminal differentiation

To understand the developmental origins and differentiation dynamics of SPP1^+^ macrophages, we performed pseudotime trajectory analysis on the monocyte/macrophage clusters. The reconstructed developmental trajectory exhibited a branched architecture (Fig. 5A and 5B). The trajectory originated from proliferating monocytes and Ly6Chi monocytes, which first differentiated into Mertk^+^ macrophages. Subsequently, Mertk^+^ macrophages bifurcated into two distinct terminal fates: SPP1^+^ macrophages and Ltk macrophages (Fig. 5A and 5B). These results indicate that SPP1^+^ macrophages represent a terminally differentiated subset derived from circulating monocytes.

**Figure 5.**
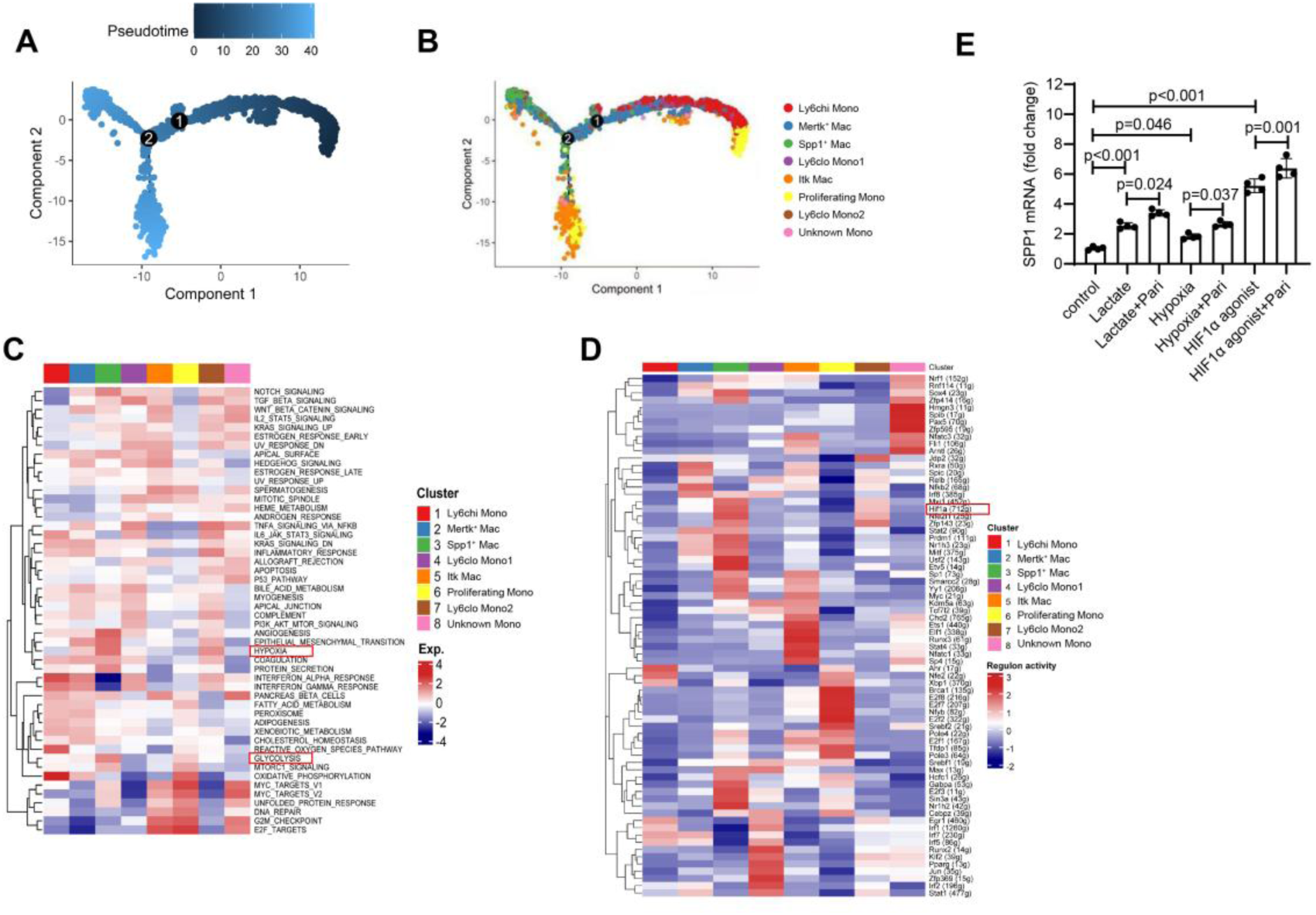
Pseudotime trajectory, pathway enrichment, and in vitro validation of SPP1^+^ macrophage differentiation. (**A**) Pseudotime mapping of hepatic macrophage/monocyte developmental trajectory. Color gradient represents pseudotime progression, with darker blue indicating earlier developmental stages and lighter blue indicating later stages along the trajectory. (**B**) Cell subset distribution along the hepatic macrophage/monocyte developmental trajectory. Each color corresponds to a unique macrophage/monocyte subset. (**C**) GSVA enrichment heatmap showing the relative activity of hallmark gene sets in each monocyte/macrophage subcluster. (**D**) Transcription factor enrichment analysis of each monocyte/macrophage subcluster. (**E**) Relative mRNA expression of SPP1 in RAW264.7 cells treated with vehicle, hypoxia, lactate (50 μM), HIF-1α agonist (Fenbendazole, 0.2 μM), with or without paricalcitol (100 nM), for 24 h. Data in panel E are presented as mean ± SD. Statistical analysis was performed using ANOVA followed by Tukey’s post hoc test. This experiment was performed in triplicate.

We next performed Gene Set Variation Analysis (GSVA) to identify signaling pathways specifically enriched in SPP1^+^ macrophages. Compared with other subsets, SPP1^+^ macrophages showed significant enrichment of hypoxia and glycolysis pathways (Fig. 5C). Consistent with these pathway signatures, transcription factor enrichment analysis revealed that HIF-1α were among the most significantly enriched transcription factors in SPP1^+^ macrophages (Fig. 5D), suggesting that hypoxia, glycolysis, and the master hypoxia regulator HIF1α may play critical roles in driving monocyte-to-SPP1^+^ macrophage differentiation.

To experimentally validate these findings, we treated RAW264.7 monocyte/macrophage cells with hypoxia, lactate (a glycolysis product), or a HIF1α agonist, alone or in combination with paricalcitol. Strikingly, each of these stimuli significantly upregulated SPP1 expression, and paricalcitol synergistically enhanced their effects (Fig. 5E). Collectively, these data demonstrate that SPP1^+^ macrophages originate from circulating monocytes and undergo terminal differentiation via a trajectory involving hypoxia, glycolysis, and HIF1α signaling, and that VDR activation synergizes with these cues to promote SPP1^+^ macrophage differentiation.

### VDR activation promotes glycolytic reprogramming in macrophages through induction of pro-glycolytic genes

Given the enrichment of glycolysis pathway in SPP1^+^ macrophages, we next investigated the relationship between VDR signaling, glycolysis, and SPP1^+^ macrophage differentiation. Analysis of glycolysis scores in monocyte/macrophage subsets revealed that infection increased glycolysis in hepatic monocyte/macrophages, and paricalcitol treatment further augmented this effect (Fig. 6A). Among all subsets, SPP1^+^ macrophages exhibited the highest glycolysis score (Fig. 6B and 6C), mirroring the VD response score distribution observed earlier. We then examined the correlation between glycolysis score and VD response score across individual monocyte/macrophage cells. A mild but significant positive correlation was observed (R = 0.047, *P* = 0.0015) (Supplementary Fig. 6A). To further explore the functional link, we stratified monocyte/macrophage cells into high and low VD response score groups (Supplementary Fig. 6B). Differential gene expression analysis revealed that SPP1 was the most significantly upregulated gene in the high VD response score group (Supplementary Fig. 6C). Moreover, GSEA showed that the glycolysis pathway was significantly enriched in cells with high VD response scores (Supplementary Fig. 6D). These findings suggest that VDR activation coordinates glycolytic reprogramming and SPP1 expression in macrophages. We next validated these observations experimentally. Isolated primary hepatic macrophages from infected mice showed increased intracellular lactate levels, which were further elevated by paricalcitol treatment (Fig. 6D). *In vitro*, paricalcitol treatment of RAW264.7 cells accelerated both glucose consumption and lactate production, indicating enhanced glycolytic flux (Fig. 6E and 6F).

**Figure 6.**
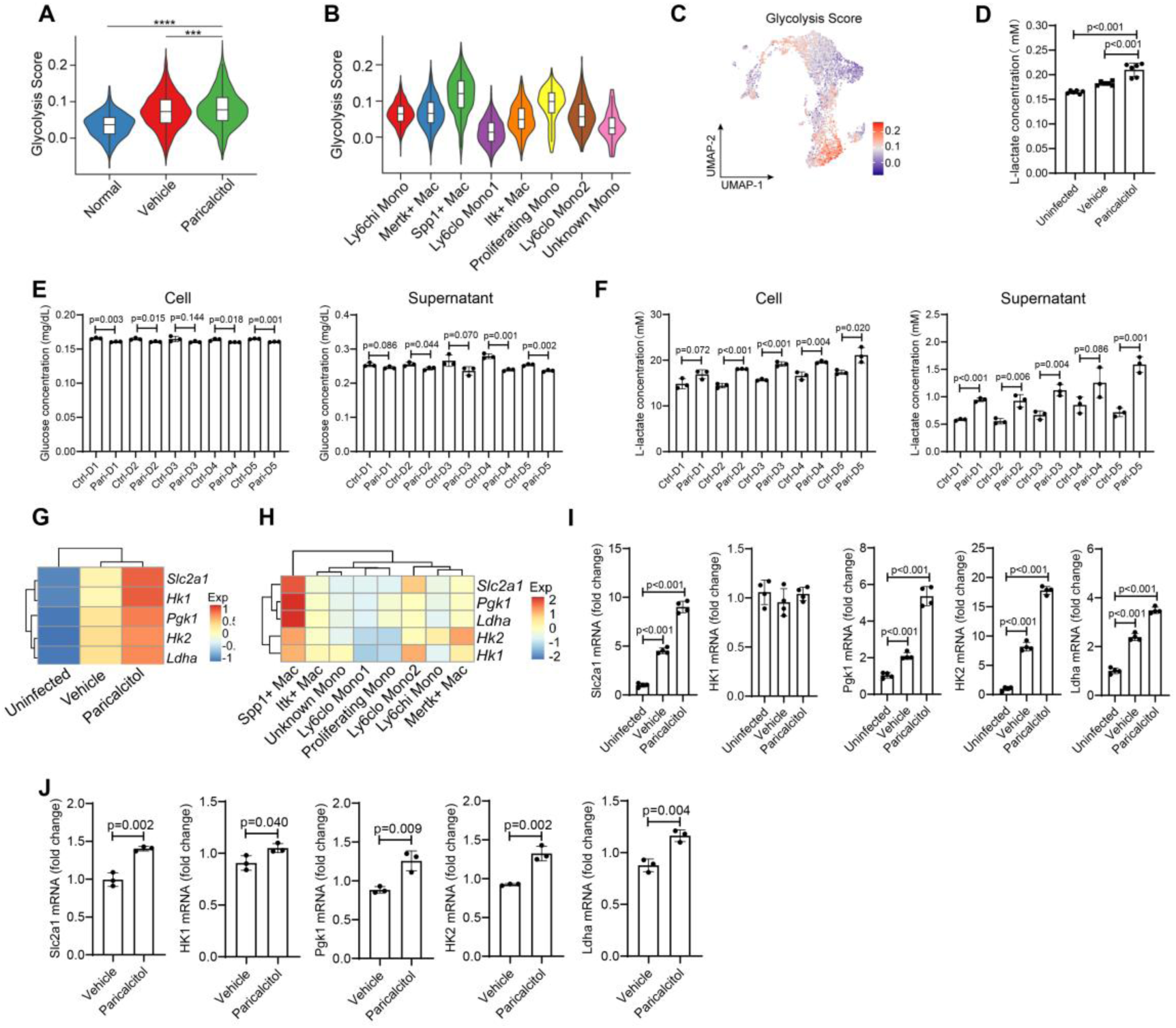
Glycolysis is enhanced in SPP1^+^ macrophages and induced by VDR activation. (**A**) Glycolysis scores in total hepatic macrophages from uninfected, infected vehicle-treated, and infected paricalcitol-treated mice. (**B**) Glycolysis scores across monocyte/macrophage subclusters. (**C**) UMAP plots of monocyte/macrophage cells colored by glycolysis score. (**D**) Intracellular lactate concentration in isolated primary hepatic macrophages from uninfected, infected vehicle-treated, and infected paricalcitol-treated mice. (**E**) Glucose concentration in the intracellular compartment and culture supernatant of RAW264.7 cells treated with vehicle or paricalcitol (100 nM) at different time points. (**F**) Lactate concentration in the intracellular compartment and culture supernatant of RAW264.7 cells treated as in (**E**). (**G**) Heatmap showing expression of glycolysis-related genes (Slc2a1, Hk1, Pgk1, Hk2, Ldha) in monocyte/macrophages across the three groups. (**H**) Heatmap showing expression of glycolysis-related genes in monocyte/macrophage subclusters. (**I**) Relative mRNA expression of glycolysis-related genes in isolated primary hepatic macrophages from uninfected, infected vehicle-treated, and infected paricalcitol-treated mice, determined by qPCR. (**J**) Relative mRNA expression of glycolysis-related genes in RAW264.7 cells treated with vehicle or paricalcitol (50 nM) for 24 h, determined by qPCR. Data from panels A and B are presented as violin plots, with embedded boxplots showing the median (center line) and interquartile range (box boundaries). The width of each violin represents the kernel density estimation of the data distribution. Statistical analysis was performed using the Kruskal-Wallis, followed by Dunn’s post-hoc test with Bonferroni correction. \*\*\*\**p*<0.0001. Data from panels D, E, F, I, and J are presented as mean ± SD. Statistical analysis was performed using ANOVA followed by Tukey’s post hoc test. These experiments were performed in triplicate.

We further analyzed the mechanisms by which the VDR pathway promotes glycolytic metabolic reprogramming in macrophages. Single-cell transcriptomic data revealed that key pro-glycolytic genes, including Slc2a1, Hk1, Pgk1, Hk2, and Ldha, were upregulated upon infection and further increased by paricalcitol (Fig. 6G). These genes were predominantly enriched in SPP1^+^ macrophages (Fig. 6H). qPCR analysis of primary hepatic macrophages and RAW264.7 cells confirmed that paricalcitol upregulates these glycolytic genes (Fig. 6I and 6J). Collectively, these data demonstrate that VDR activation promotes glycolytic reprogramming in macrophages by inducing the expression of key pro-glycolytic genes.

### Lactate and HIF1α synergistically induce VDR expression in macrophages by binding to the *Vdr* promoter and enhancing histone lactylation

Given the reciprocal relationship between VDR activation and glycolysis, we next investigated whether glycolysis could regulate VDR expression. Single-cell RNA sequencing data revealed that VDR expression in hepatic macrophages was upregulated upon infection and further increased by paricalcitol treatment (Fig. 7A). qPCR analysis of isolated primary hepatic macrophages confirmed these findings (Fig. 7B). We hypothesized that lactate, a glycolytic metabolite, and HIF1α, a master transcription factor activated under hypoxia, might drive VDR expression. *In vitro*, treatment of RAW264.7 cells with either lactate or a HIF1α agonist significantly induced VDR mRNA expression, and the combination produced a marked synergistic effect (Fig. 7C). Analysis of the mouse *Vdr* gene promoter identified two high-scoring HIF1α binding sites (Fig. 7D). Dual-luciferase reporter assays confirmed that these sites are functional (Fig. 7E). Cleavage under targets and tagmentation (Cut&Tag)-qPCR demonstrated that HIF1α agonist stimulation enhanced HIF1α enrichment at the *Vdr* promoter (Fig. 7F), while lactate stimulation increased histone lactylation at the same region (Fig. 7G). Notably, combining HIF1α agonist and lactate synergistically boosted HIF1α recruitment to the *Vdr* promoter (Fig. 7H). *In vivo*, infection increased both HIF1α binding and histone lactylation at the *Vdr* promoter in hepatic macrophages, and paricalcitol further augmented these modifications (Fig. 7I). Collectively, these data reveal a positive feedback loop whereby glycolytic lactate and HIF1α cooperatively induce VDR expression in macrophages during schistosomiasis, potentially amplifying the antifibrotic VDR response.

**Figure 7.**
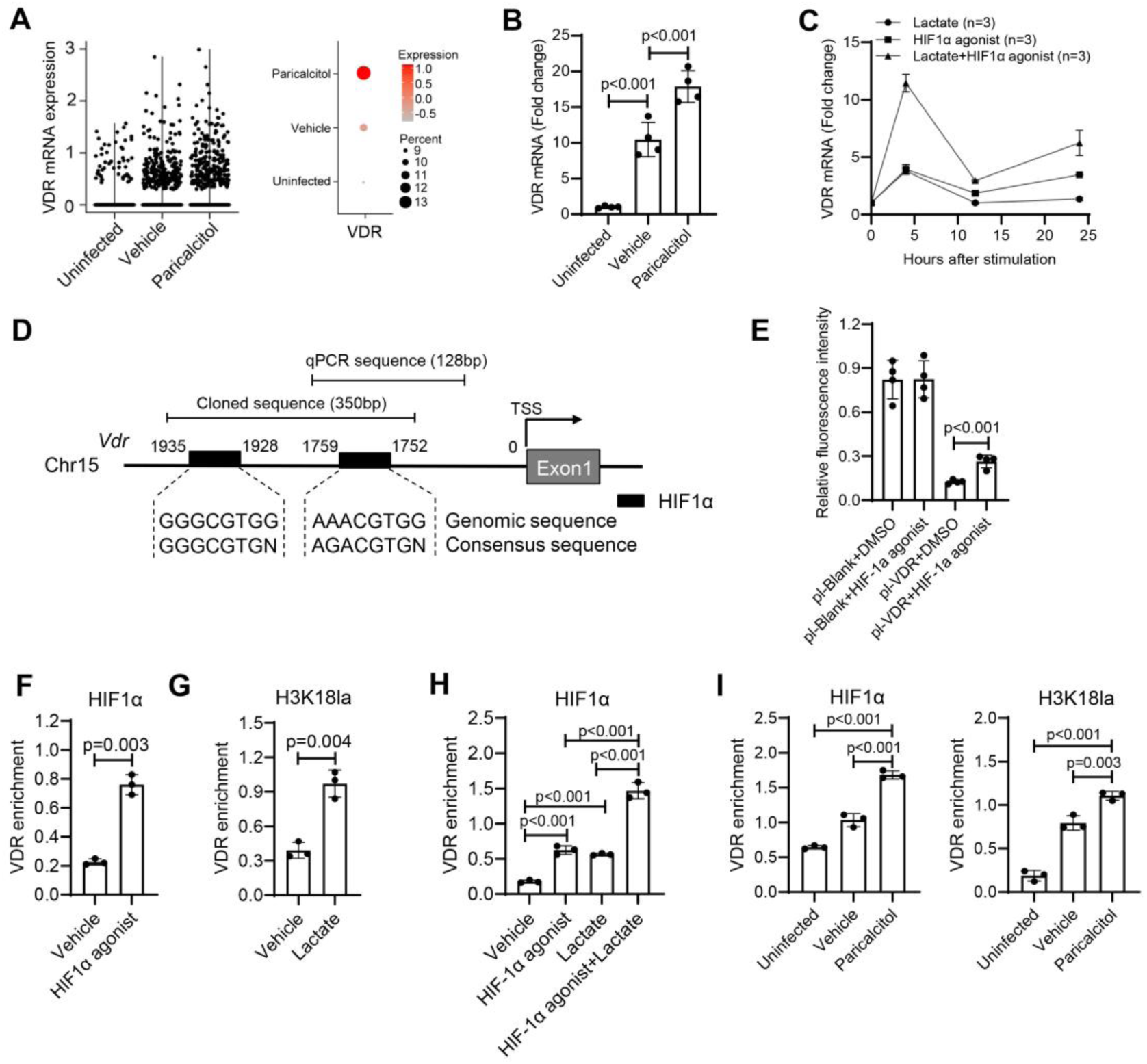
HIF1α and lactate synergistically induce VDR expression in macrophages by binding to the *Vdr* promoter and promoting histone lactylation. (**A**) Scatter plot (left) and dot plot (right) showing VDR mRNA expression across uninfected, vehicle-treated, and paricalcitol-treated groups, analyzed by single-cell transcriptomic sequencing. (**B**) Relative mRNA expression of VDR mRNA in isolated primary hepatic macrophages from the three groups, determined by qPCR. (**C**) Time-course analysis of VDR mRNA expression in RAW264.7 cells treated with control, lactate (50 μM), HIF1α agonist (Fenbendazole, 0.2 μM), or lactate plus HIF1α. (**D**) Schematic of the mouse *Vdr* gene promoter region showing two predicted high-score HIF1α binding sites. The diagram also depicts the 350-bp cloned sequence spanning both HIF1α binding sites, which was utilized for dual-luciferase reporter assays to validate transcriptional regulation, and the 128-bp qPCR sequence, which was targeted for Cut&Tag-qPCR experiments to assess HIF1α DNA binding and histone lactylation modifications. (**E**) Dual-luciferase reporter assay in RAW264.7 cells transfected with blank or *Vdr* promoter constructs, treated with HIF1α agonist for 24 h. (**F**) Cut&Tag-qPCR analysis of HIF1α enrichment at the *Vdr* promoter in RAW264.7 cells treated with HIF1α agonist. (**G**) Cut&Tag-qPCR analysis of H3K18la (histone lactylation) enrichment at the *Vdr* promoter in RAW264.7 cells treated with lactate. (**H**) Cut&Tag-qPCR analysis of HIF1α enrichment at the *Vdr* promoter in RAW264.7 cells treated with lactate, HIF1α agonist, or both. (**I**) Cut&Tag-qPCR analysis of HIF1α and H3K18la enrichment at the *Vdr* promoter in isolated primary hepatic macrophages from uninfected, infected vehicle-treated, and infected paricalcitol-treated mice. Data from panels B, C, and E-I are presented as mean ± SD. Comparisons between two groups were performed using two-tailed Student’s t-test. Multiple group comparisons were performed by one-way ANOVA followed by Tukey’s post hoc test. These experiments were performed in triplicate.

## DISCUSSION

The prevailing paradigm of VDR-mediated antifibrosis has largely focused on its direct actions on HSCs, where liganded VDR antagonizes TGF-β/SMAD signaling (*8*). Consistent with this model, our previous work in schistosomiasis showed that VDR expression in HSCs is negatively regulated by the IFN-γ/IRF2/miR-351 axis, and that VDR activation directly inhibits HSC activation (*9*). However, the present study uncovers a previously unrecognized and functionally dominant role of VDR in macrophages. We found that VDR expression is highest in hepatic macrophages among liver-resident cells, and that monocytes/macrophages display the highest vitamin D response signature upon paricalcitol treatment. More importantly, myeloid-specific *Vdr* knockout mice developed exacerbated liver fibrosis, directly demonstrating that macrophage VDR is essential for antifibrosis. Strikingly, paricalcitol treatment in myeloid-specific *Vdr* knockout mice still partially reduced fibrotic gene expression and hydroxyproline content, likely reflecting residual VDR activity in HSCs or other cell types, but its beneficial effects on granuloma formation and liver injury were completely abolished. This dissociation indicates that while HSC-intrinsic VDR signaling may directly suppress fibrotic gene expression, the anti-inflammatory and hepatoprotective effects of VDR activation critically depend on macrophage VDR signaling. Our findings are strongly supported by recent studies showing that VDR activation in macrophages ameliorates hepatic inflammation, steatosis, and insulin resistance, and protects against endoplasmic reticulum stress-induced liver injury (*13*,*14*). Collectively, these observations shift the paradigm from a HSC-centric view to a more integrated model in which macrophage VDR acts as a central rheostat controlling the inflammatory microenvironment and hepatocyte integrity, thereby limiting schistosomiasis-induced hepatic fibrosis. This expanded understanding opens new avenues for cell-type-specific therapeutic strategies targeting macrophage VDR signaling in fibrotic liver diseases.

The functional role of SPP1⁺ macrophages in fibrotic diseases remains a subject of considerable debate. On one hand, numerous studies have implicated SPP1⁺ macrophages as pro-fibrotic effectors across multiple organs. In liver cirrhosis, scar-associated macrophages characterized by high SPP1 expression are enriched in fibrotic niches and promote tissue fibrosis and angiogenesis (*15*). Similar pro-fibrotic populations have been identified in idiopathic pulmonary fibrosis, where SPP1⁺ macrophages accumulate in fibrotic lung regions and correlate with disease severity (*16*). Notably, SPP1⁺ macrophages exhibit cross-organ conservation in fibrotic diseases, making them attractive therapeutic targets (*17*,*18*). On the other hand, emerging evidence supports a protective, anti-fibrotic function for macrophage-derived SPP1 in certain contexts. Han and colleagues demonstrated that macrophage-derived SPP1 (osteopontin) protects against nonalcoholic steatohepatitis by inducing oncostatin-M expression in macrophages, which in turn upregulates arginase-2 in hepatocytes, enhances fatty acid oxidation, and reduces steatosis and fibrosis (19). In a completely different setting, Dolci *et al*. showed that tumor-associated macrophages with high SPP1 expression promote spinal cord repair and inhibit post-injury fibrosis, an effect dependent on SPP1 secretion (*20*). Our study firmly supports the protective arm of this dichotomy. We found that SPP1⁺ macrophages are the critical effector cells downstream of VDR activation, and that selective depletion of SPP1⁺ macrophages using SPP1-DTR mice significantly aggravated schistosomiasis-induced hepatic fibrosis. Nevertheless, the precise molecular mechanism by which SPP1⁺ macrophages limit fibrosis remains incompletely understood and is a priority for future investigation. Collectively, these seemingly contradictory findings suggest that the function of SPP1⁺ macrophages is highly context-dependent, shaped by disease etiology, pathological stage, and local microenvironmental cues. Therefore, therapeutic strategies targeting SPP1⁺ macrophages must be carefully tailored to the specific disease context.

Consistent with previous studies, our data indicate that hypoxia-glycolysis axis is a central driver of SPP1⁺ macrophage differentiation (*21*,*22*). Moreover, VDR activation synergizes with hypoxia, lactate, and HIF1α to promote this macrophage phenotype, although the precise molecular mechanism remains to be fully elucidated. Importantly, we have uncovered a lactate-HIF1α–VDR positive feedback circuit: glycolysis generate lactate, which together with HIF1α induces VDR transcription; activated VDR in turn promotes further glycolytic reprogramming, producing more lactate, thereby amplifying the protective response. This feedback loop aligns with a growing body of literature demonstrating that acute hypoxia and metabolic reprogramming are beneficial for tissue repair (*23*). In models of myocardial infarction, ischemic preconditioning, and colitis, moderate hypoxia and lactate administration limit inflammation and promote regeneration, effects that are at least partly dependent on VDR (*24-26*). In the context of schistosomiasis, the egg granuloma itself is a hypoxic niche (*27*). Our data suggest that this hostile microenvironment paradoxically triggers a self-limiting protective circuit: local lactate and HIF1α upregulate VDR in infiltrating monocytes/macrophages, and VDR activation in turn drives their differentiation into SPP1⁺ macrophages, which limit excessive fibrosis. Thus, this endogenous metabolic-epigenetic circuit represents a self-reinforcing defense mechanism and suggests that targeting the lactate–HIF1α–VDR axis may offer a new therapeutic strategy for fibrotic diseases.

In summary, this study redefines the cellular and metabolic basis of VDR-mediated protection against schistosomiasis-induced hepatic fibrosis. We demonstrate that macrophage VDR is the primary driver of the antifibrotic response, acting by promoting the differentiation of a protective SPP1⁺ macrophage subset. Mechanistically, we uncover a positive feedback circuit in which lactate and HIF1α cooperatively induce VDR expression, and VDR activation in turn enhances glycolysis and SPP1⁺ macrophage differentiation, creating a self-amplifying loop that limits fibrogenesis (Supplementary Fig. 7). These findings resolve the controversial role of SPP1⁺ macrophages in fibrosis by establishing their protective function in this context, and they shift the paradigm of VDR action from a HSC-centric model to an integrated immunometabolic circuit centered on macrophages. Collectively, our work identifies the lactate-HIF1α-VDR-SPP1 axis as a novel endogenous defense mechanism and provides a strong rationale for targeting this pathway, particularly by VDR agonists, as a promising therapeutic strategy for schistosomiasis-associated liver fibrosis and potentially other fibrotic liver diseases.

## MATERIALS AND METHODS

### Ethics statement

All animal experiments were performed in strict accordance with the Guide for the Care and Use of Laboratory Animals of the National Institutes of Health, and were approved by the Animal Ethics Committee of Naval Medical University (approval number: 2021-0006). All procedures were performed under carbon dioxide anesthesia, and every effort was made to minimize animal suffering.

### Animals and parasite infection

Six-week-old male C57BL/6J mice were purchased from the Experimental Animal Center of Naval Medical University. VDR^flox/flox^ (VDR^fl/fl^) mice (Cat. NO. NM-CKO-210040) and SPP1^LSL-DTR-EGFP^ mice (Cat. NO. NM-KI-234008) mice were obtained from Shanghai Model Organisms Center, Inc.. Lyz2-Cre mice were obtained from Jackson Laboratory (stock #004781). Mice were housed under specific pathogen-free conditions with autoclaved food and water ad libitum. *Schistosoma japonicum* (*S. japonicum*) (Chinese mainland strain) cercariae were obtained from infected *Oncomelania hupensis* snails supplied by the Shanghai Veterinary Research Institute, Chinese Academy of Agricultural Sciences. For infection, mice were percutaneously exposed to 20 cercariae through the abdominal skin as described previously.

### Drug treatment

Paricalcitol (Hengrui Pharmaceutical Co., Ltd., Jiangsu, China; commercial injection formulation) was diluted in normal saline containing 5% propylene glycol. Infected mice received paricalcitol (4 µg/kg body weight) or vehicle (normal saline containing 5% propylene glycol) by intraperitoneal injection three times per week, starting immediately after infection until sacrifice. For *in vitro* experiments, paricalcitol was used at a final concentration of 100 nM.

### Tissue processing and histopathology

Livers were harvested at indicated time points, fixed in 4% paraformaldehyde, and embedded in paraffin. Sections (4 µm) were stained with hematoxylin and eosin (H&E) for granuloma size measurement or with Masson’s trichrome for collagen deposition. Granuloma size was quantified by measuring at least 10 randomly selected granulomas per section. Fibrosis was scored by multiplying the blue density (scale 1–4) and the area involved (scale 1–4) for each granuloma, giving a maximum score of 16 (*28*). This study adopted a double-blind design, in which the pathologists performing the pathological examination were unaware of the group assignments.

Hydroxyproline content, as a biochemical indicator of total collagen, was measured using a colorimetric assay kit (Nanjing Jiancheng Bioengineering Institute, Nanjing, China) according to the manufacturer’s protocol. Briefly, 50 mg of liver tissue was hydrolyzed in 6 M HCl at 110 °C overnight, and the hydroxyproline concentration was calculated from a standard curve.

### Isolation of primary liver cells

Primary hepatocytes, hepatic stellate cells (HSCs), liver endothelial sinusoidal cells (LESCs), and hepatic macrophages were isolated from mouse livers by a two-step collagenase perfusion method followed by density gradient centrifugation and immunomagnetic purification as previously described (*29*). In brief, the liver was perfused in situ via the portal vein with Ca²⁺/Mg²⁺-free Hanks’ balanced salt solution (HBSS) followed by 0.05% collagenase IV and 0.02% pronase E (Sigma-Aldrich, St. Louis, MO, USA) in HBSS containing Ca²⁺ and Mg²⁺. The digested liver was gently disrupted, and the cell suspension was filtered through a 70 µm cell strainer. Hepatocytes were pelleted by low-speed centrifugation (50 × g, 4 min). The non-parenchymal cell fraction was further separated on an 11.5% iodixanol (OptiPrep; Axis-Shield, Oslo, Norway) gradient. HSCs were collected from the top layer and further purified by negative selection using CD45 magnetic beads (Miltenyi Biotec, Auburn, CA, USA). Hepatic macrophages were isolated by positive selection from the non-parenchymal cell fraction using F4/80 microbeads. LESCs were isolated by positive selection from the non-parenchymal cell fraction using CD146 microbeads.

### RNA extraction and quantitative real-time PCR (qPCR)

Total RNA was extracted from tissues or cells using TRIzol reagent (Invitrogen, Carlsbad, CA, USA) according to the manufacturer’s instructions. cDNA was synthesized with the PrimeScript RT reagent kit (Takara, Kusatsu, Japan). qPCR was carried out using SYBR Green Master Mix (Roche, Basel, Switzerland) on a LightCycler 480 system (Roche). The relative expression was calculated by the 2^⁻ΔΔCt^ method using GAPDH as internal controls (*30*). Primer sequences are listed in Supplementary Table S1.

### Western blotting

Cells or tissues were lysed in RIPA buffer (Beyotime, Shanghai, China) supplemented with protease and phosphatase inhibitor cocktails (Thermo Fisher Scientific, Waltham, MA, USA). Protein concentrations were determined by the BCA method (Pierce, Rockford, IL, USA). Equal amounts of protein (20-30 µg) were separated by 10–12% SDS-PAGE and transferred onto nitrocellulose membranes (PALL, Port Washington, NY, USA). After blocking with 5% non-fat milk in TBST, membranes were incubated overnight at 4 °C with primary antibodies against VDR (CST, 12550) or GAPDH (Abcam, ab9485). IRDye 800CW secondary antibodies (LI-COR, Lincoln, NE, USA) were used for detection with an Odyssey infrared imaging system.

### Single-cell RNA sequencing (scRNA-seq)

Hepatic CD45⁺ immune cells were isolated from uninfected, infected vehicle-treated, and infected paricalcitol-treated mice at day 42 post-infection by fluorescence-activated cell sorting (FACS). Cell viability (>90%) was assessed by trypan blue exclusion. scRNA-seq libraries were prepared using the Chromium Single Cell kit (v3.1, 10× Genomics, Pleasanton, CA, USA) following the manufacturer’s protocol, targeting ∼10000 cells per sample. Libraries were sequenced on an Illumina NovaSeq 6000 platform with a paired-end 150 bp read length.

Raw sequencing data were processed with Cell Ranger (v6.0, 10× Genomics) using the mouse reference genome mm10. Cells with <200 or >2500 detected genes and >20% mitochondrial reads were excluded. Downstream analysis was performed with Seurat (v4.3) in R (v4.5). Principal component analysis (PCA) was conducted on highly variable genes, and the top 30 PCs were used for uniform manifold approximation and projection (UMAP) dimensionality reduction and graph-based clustering (resolution = 0.2). Clusters were annotated by canonical marker genes from the literature. For monocyte/macrophage subclustering, cells annotated as monocytes/macrophages were extracted and re-clustered with a resolution of 0.2.

A vitamin D (VD) response score was calculated for each cell using the AddModuleScore function in Seurat based on a list of known VDR target genes (e.g., VDR, Cyp27b1) compiled from Gene Ontology and previous reports. Glycolysis scores were computed using the same method with a curated set of glycolysis-related genes (e.g., Slc2a1, Hk1, Hk2, Pgk1, Ldha). Gene set variation analysis (GSVA) was performed using the GSVA package with the MSigDB hallmark gene sets. Transcription factor enrichment analysis was conducted using the Single-Cell Regulatory Network Inference and Clustering (SCENIC) pipeline (*31*). Pseudotime trajectory analysis was performed using Monocle2 with the DDRTree method, using the eight monocyte/macrophage subclusters as input.

Raw single-cell transcriptomic sequencing data have been deposited in the Genome Sequence Archive (GSA) of the National Genomics Data Center (NGDC), China National Center for Bioinformation (CNCB), under accession number CRA044939, which can be accessed at https://ngdc.cncb.ac.cn/gsa/browse/CRA044939.

### Generation of myeloid-specific VDR knockout mice

To generate macrophage-specific VDR knockout, VDR^fl/fl^ mice were crossed with Lyz2-Cre transgenic mice. The resulting VDR^fl/fl^;Lyz2-Cre⁺/⁻ (designated VDR^lyz2-Cre^) mice and littermate VDR^fl/fl^;Lyz2-Cre⁻/⁻ (control) mice were genotyped by PCR using tail DNA. VDR deletion in macrophages was confirmed by western blot of F4/80-purified hepatic macrophages.

### Generation of SPP1^+^ macrophage reporter/depletion mice

SPP1^LSL-DTR-EGFP^ mice were generated by inserting a loxp-stop-loxp (LSL) cassette followed by a DTR-2A-EGFP sequence into the *SPP1* locus using CRISPR/Cas9-mediated homologous recombination. These mice were crossed with Lyz2-Cre mice to obtain SPP1^LSL-DTR-EGFP/+^;Lyz2-Cre⁺/⁻ (designated SPP1-DTR) mice, in which Cre-mediated excision removes the stop cassette, allowing DTR and EGFP expression specifically in SPP1⁺ macrophages. Control mice were SPP1^LSL-DTR-EGFP/+^;Lyz2-Cre⁻/⁻ littermates. For depletion, mice received intraperitoneal injections of diphtheria toxin (DT, 10 ng/g body weight, Sigma-Aldrich) or PBS every other day starting at day 21 post-infection. Depletion efficiency was assessed by flow cytometry for EGFP⁺ cells among CD45⁺F4/80⁺ macrophages.

### Cell culture and treatments

RAW264.7 murine macrophage cells (ATCC, TIB-71) were maintained in DMEM (HyClone, Logan, UT, USA) supplemented with 10% fetal bovine serum (Gibco), 100 U/mL penicillin, and 100 μg/mL streptomycin at 37 °C in a 5% CO₂ humidified incubator. For hypoxia treatment, cells were placed in a hypoxia chamber (1% O₂, 5% CO₂, 94% N₂) for 24 h. For lactate stimulation, sodium L-lactate (MCE) was added to the culture medium at a final concentration of 50 μM. HIF-1α agonist Fenbendazole (0.2 μM, MCE) was used to stabilize HIF-1α. Glucose and lactate concentrations in the cell compartment and culture supernatant were measured using commercial kits (Beyotime Biotechnology, China) according to the manufacturer’s instructions.

### Dual-luciferase reporter assay

A 350-bp fragment of the mouse *Vdr* promoter region containing two predicted HIF-1α binding sites was amplified from mouse genomic DNA and cloned into the pGL3-Basic vector (Promega, Madison, WI, USA). RAW264.7 cells (1 × 10⁵ cells/well in 24-well plates) were co-transfected with 0.5 μg of reporter plasmid, 20 ng of pRL-TK Renilla luciferase control vector, and 0.2 μM HIF-1α agonist or vehicle, using Lipofectamine 3000 (Invitrogen). After 24 h, firefly and Renilla luciferase activities were measured using the Dual-Luciferase Reporter Assay System (Promega) on a GloMax 20/20 luminometer. Relative luciferase activity was calculated as firefly/Renilla ratio.

### Cut&Tag assay

The cleavage under targets and tagmentation (CUT&Tag) assay was performed using the Hyperactive Universal CUT&Tag Assay Kit for Illumina (TD903; Vazyme Biotech) following the manufacturer’s protocol. Briefly, RAW264.7 cells or freshly isolated primary hepatic macrophages (1 × 10⁵ cells per reaction) were harvested, bound to concanavalin A beads, and incubated with primary antibodies against HIF-1α (CST, 36169) or H3K18la (PTM-Bio, Hangzhou, China) overnight at 4 °C. After washing, cells were incubated with guinea pig anti-rabbit IgG secondary antibody, followed by pAG-Tn5 transposase adapter complex. Targeted DNA fragments were amplified by PCR using primers flanking the *Vdr* promoter region. Libraries were purified and quantified by qPCR. Enrichment was calculated relative to input chromatin.

### Statistical analysis

All data are presented as mean ± standard deviation (SD). Comparisons between two groups were performed using two-tailed Student’s *t*-test. Multiple group comparisons were performed by one-way ANOVA followed by Tukey’s post hoc test. For scRNA-seq data, differential expression between clusters was assessed using the Kruskal-Wallis, followed by Dunn’s post-hoc test with Bonferroni correction. Correlation between vitamin D (VD) response score and glycolysis score was analyzed by Pearson correlation. A *P*-value < 0.05 was considered statistically significant. All statistical analyses were performed using SPSS (version 26.0, IBM Corp., Armonk, NY, USA) or R software (v4.5).

## Acknowledgments

We thank Professor Zhiqiang Fu from the Shanghai Veterinary Research Institute, Chinese Academy of Agricultural Sciences, for providing S. japonicum cercariae and assisting with mouse infection. We are grateful to Professor Weiqing Pan from Naval Medical University for valuable advice on the research and manuscript preparation.

## Funding

This work was supported by grants from the National Natural Science Foundation of China: 82322061 (X.H.), 82173640 (X.H.), 82404398 (X.F.).

## Author contributions

Conceptualization: X.G., X.F., and X.H. Methodology: Z.X., P.Y., H.X., R.Y., B.L., and F.Z. Software: Z.X., and X.F. Validation: X.F., and X.H. Formal analysis: Z.X., P.Y., and H.X. Investigation Resources: X.G., X.F., and X.H. Data curation: X.G., X.F., and X.H. Writing: X.H. Visualization: Z.X., P.Y., X.F., and H.X. Supervision: H.X. Funding acquisition: X.F., and X.H.

## Competing interests

The authors declare that they have no competing interests.

## Data, code, and materials availability

All data and code needed to evaluate and reproduce the results in the paper are present in the paper and/or the Supplementary Materials. All materials used in this study are commercially available. Raw single-cell transcriptomic sequencing data have been deposited in the Genome Sequence Archive (GSA) of the National Genomics Data Center (NGDC), China National Center for Bioinformation (CNCB), under accession number CRA044939, which can be accessed at https://ngdc.cncb.ac.cn/gsa/browse/CRA044939.

## Notes

### Competing Interest Statement

The authors have declared no competing interest.

