## supplementary materials for "A lactate-HIF1α-VDR positive feedback loop drives protective SPP1⁺ macrophage differentiation to inhibit schistosomiasis-induced liver fibrosis"

### **Affiliations**

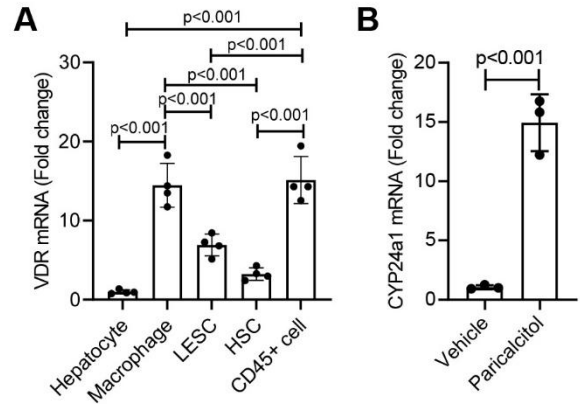

**Fig. S1. VDR is highly expressed in hepatic macrophages and functionally responsive to paricalcitol.** (A) Relative mRNA expression of VDR in freshly isolated primary hepatocytes, macrophages, hepatic stellate cells (HSCs), liver endothelial sinusoidal cells (LESCs), and CD45<sup>+</sup> immune cells from mouse liver. (B) Relative mRNA expression of CYP24A1 in primary mouse liver macrophages treated with vehicle or paricalcitol (100 nM) for 24 h. Data are presented as mean ± SD. Multiple group comparisons were performed by one-way ANOVA followed by Tukey's post hoc test. Comparisons between two groups were performed using two-tailed Student's *t*-test. The experiments were performed in triplicate.

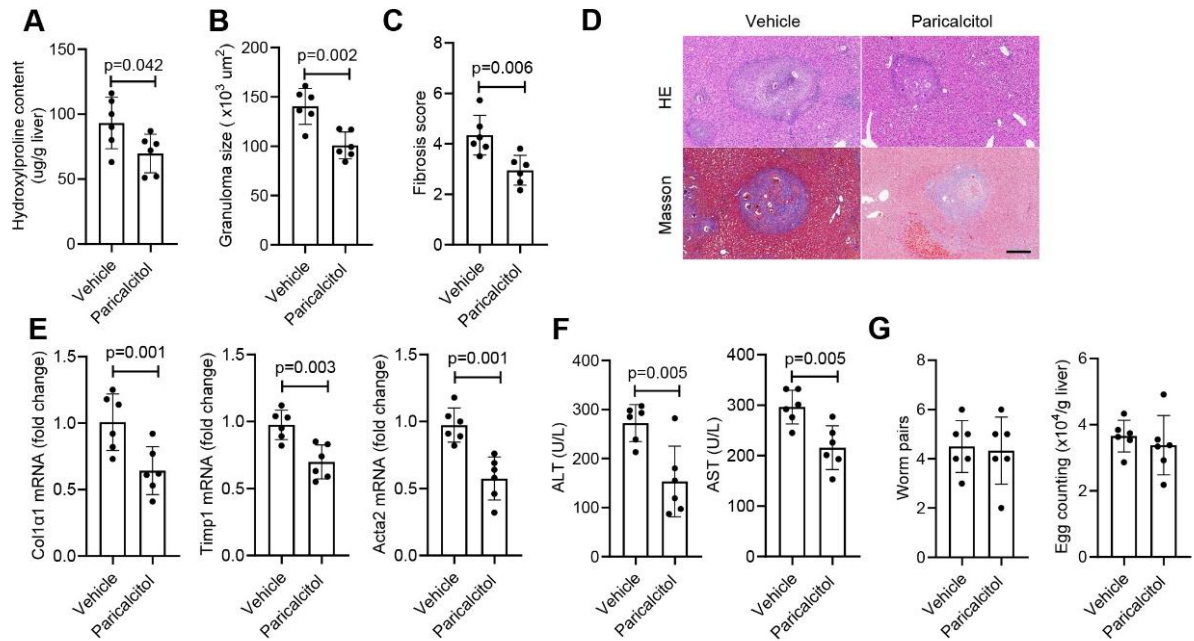

**Fig. S2. Paricalcitol treatment alleviates schistosomiasis-induced hepatic fibrosis.** C57BL/6 mice were infected with *S. japonicum* cercariae and treated with vehicle or paricalcitol three times per week. Livers and serum were collected at day 42 post-infection. (A) Hepatic hydroxyproline content. (B) Granuloma size measured from H&E-stained liver sections. (C) Fibrosis score assessed by Masson's trichrome staining. (D) Representative images of Masson's trichrome staining (upper) and H&E staining (lower) of liver sections. Scale bar, 200 $\mu\text{m}$. (E) Relative mRNA expression of fibrotic markers Col1a1, Timp1, and Acta2 in liver tissues determined by qPCR. (F) Serum ALT and AST levels. (G) Adult worm pairs and hepatic egg burden. Data are presented as mean ± SD. Comparisons between two groups were performed using two-tailed Student's *t*-test. The experiments were performed in triplicate.

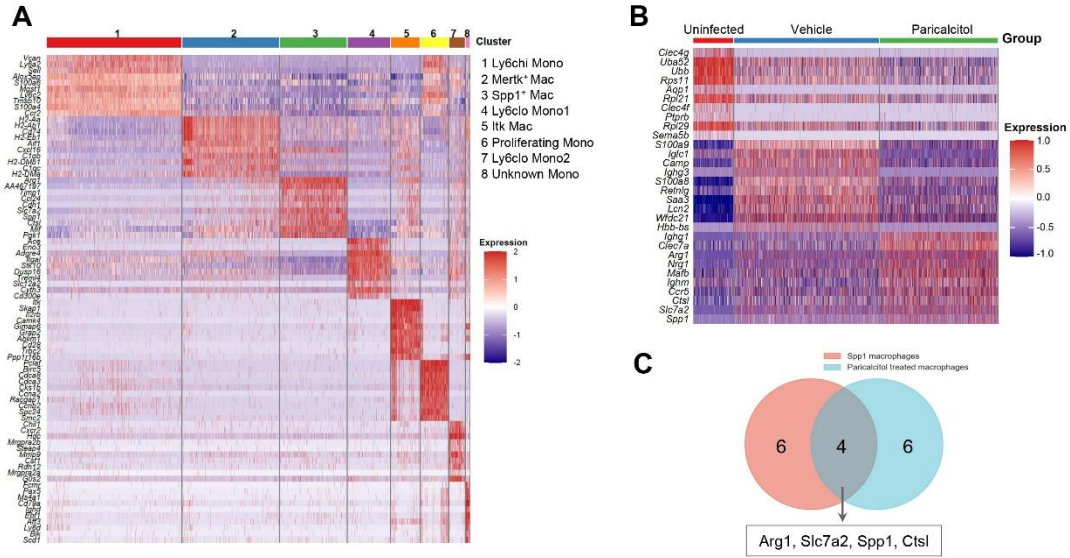

**Fig. S3. Signature gene sets of monocyte/macrophage subclusters.** (A) Heatmap showing the expression of top 10 marker genes used to annotate the eight monocyte/macrophage subclusters (related to Fig. 2A). Each column represents an individual cell, and each row represents a gene. Color scale indicates normalized expression levels. (B) Heatmap showing the expression of top 10 marker genes in macrophages from uninfected, infected vehicle-treated, and infected paricalcitol-treated groups. (C) Venn diagram illustrating the overlap between marker genes expressed in SPP1<sup>+</sup> macrophages and marker genes expressed in paricalcitol-treated macrophages. The four overlapping genes (Arg1, Slc7a2, Spp1, and Ctsl) were used as the signature gene set for SPP1<sup>+</sup> macrophages.

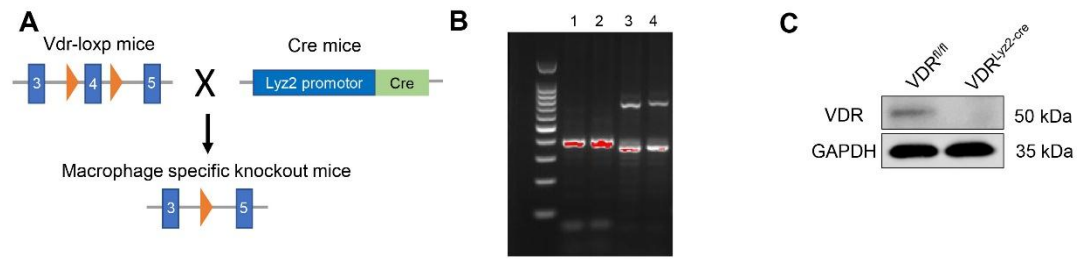

**Fig. S4. Generation and validation of myeloid-specific VDR knockout mice.** (A) Schematic diagram of the targeting strategy for generating *VDR<sup>fl/fl</sup>* mice and crossing with *Lyz2-Cre* mice to produce myeloid-specific VDR knockout (*VDR<sup>lyz2-Cre</sup>*) mice. (B) Genotyping PCR analysis of tail DNA from *VDR<sup>fl/fl</sup>* and *VDR<sup>lyz2-Cre</sup>* mice. The *VDR* floxed allele yields a 377 bp product (lane 1 and 2). The *Cre* transgene yields a 700 bp product, and the wild type (wt) allele yields a 350 bp product (lane 3 and 4). (C) Western blot analysis of VDR protein expression in isolated hepatic macrophages from *VDR<sup>fl/fl</sup>* and *VDR<sup>lyz2-Cre</sup>* mice. GAPDH was used as a loading control.

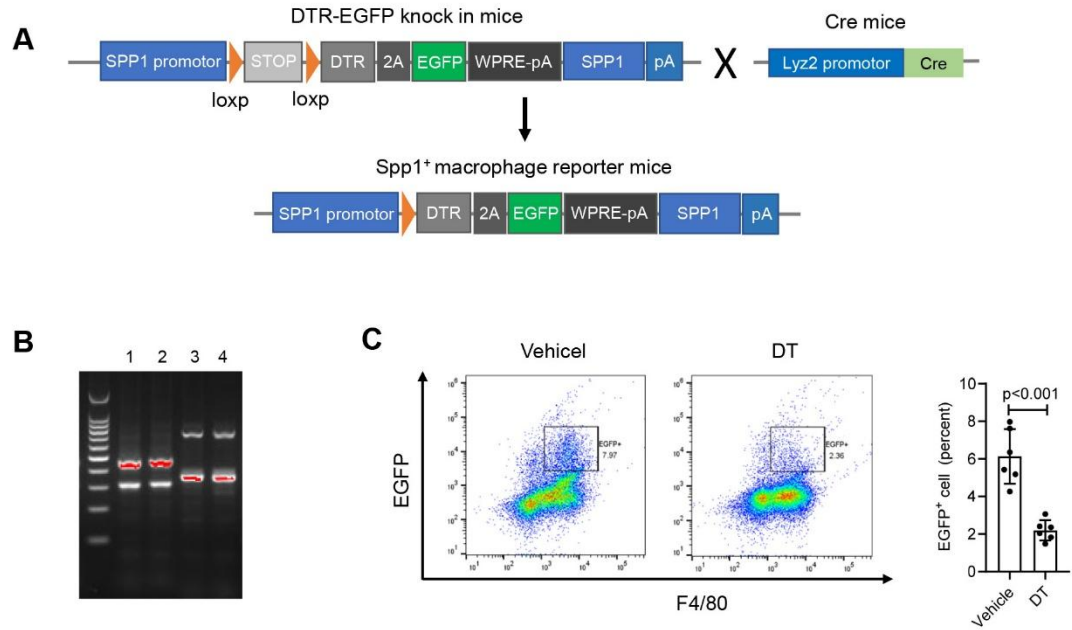

**Fig. S5. Generation and validation of SPP1<sup>+</sup> macrophage reporter/depletion mice.** (A) Schematic diagram of the targeting strategy for generating SPP1<sup>LSL-DTR-EGFP</sup> mice and crossing with Lyz2-Cre mice to produce SPP1-specific DTR-EGFP reporter/depletion mice. (B) Genotyping PCR analysis of tail DNA from SPP1<sup>LSL-DTR-EGFP</sup>/Lyz2-Cre<sup>+/-</sup> mice. The SPP1-DTR wild-type allele yields a 313 bp product, and the SPP1-DTR targeted allele yields a 377 bp product (lane 1 and 2). The Cre transgene yields a 700 bp product, and the Cre wild type (wt) allele yields a 350 bp product (lane 3 and 4). (C) Flow cytometry analysis of EGFP<sup>+</sup> cells among hepatic CD45<sup>+</sup> immune cells in infected SPP1-DTR mice treated with PBS or DT.

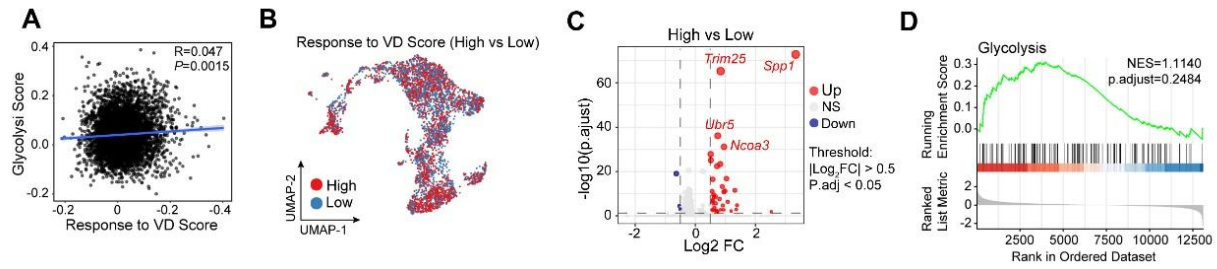

**Fig. S6. Glycolysis score positively correlates with VD response score in hepatic monocytes/macrophages.** (A) Scatter plot showing correlation between glycolysis score and VD response score in individual monocyte/macrophage cells from scRNA-seq data. Each dot represents a single cell. (B) UMAP plot of monocyte/macrophage cells colored by VD response score (high vs. low). (C) Volcano plot of differentially expressed genes between high and low VD response score groups. (D) GSEA enrichment plot showing glycolysis pathway enrichment in the high VD response score group.

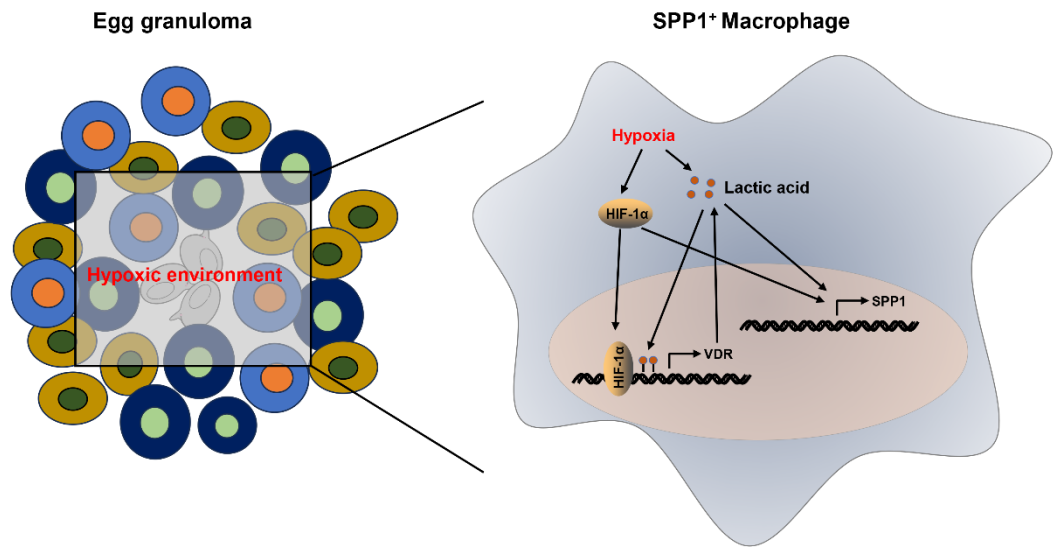

**Fig. S7. A lactate–HIF1α–VDR positive feedback circuit drives SPP1<sup>+</sup> macrophage differentiation to inhibit schistosomiasis-induced hepatic fibrosis.** Schematic illustration of the molecular mechanism by which VDR signaling in macrophages promotes SPP1<sup>+</sup> macrophage differentiation and limits schistosomiasis-induced hepatic fibrosis. The egg granuloma, the fundamental pathological feature of hepatic schistosomiasis, creates a hypoxic microenvironment. Hypoxia induces glycolytic reprogramming in infiltrating macrophages, generating lactate. Lactate drives histone H3K18 lactylation (H3K18la) at the *Vdr* promoter, while hypoxia activates HIF1α, which binds to hypoxia-response elements in the *Vdr* promoter. Lactate and HIF1α synergistically induce VDR expression in macrophages. Activated VDR further promotes glycolytic reprogramming, producing more lactate and sustaining HIF1α activity, thereby establishing a positive feedback loop that amplifies the protective response. Lactate and HIF1α, together with VDR activation, promote SPP1 expression and drive macrophage differentiation into the protective SPP1<sup>+</sup> phenotype. These SPP1<sup>+</sup> macrophages limit granulomatous inflammation and hepatic fibrosis, preserving hepatocyte integrity. Thus, the lactate–HIF1α–VDR–SPP1 axis represents an endogenous metabolic-epigenetic defense mechanism against schistosomiasis-induced liver fibrosis.

**Table S1. Primer sequences used in this study.**

| gene |  | primer sequence (5'→3') |
| --- | --- | --- |
| Hk1 | forward primer | CGGAATGGGGAGCCTTTGG |
|  | reverse primer | GCCTTCCTTATCCGTTTCAATGG |
| Slc2a1 | forward primer | CAGTTCGGCTATAACACTGGTG |
|  | reverse primer | GCCCCCGACAGAGAAGATG |
| Pgk1 | forward primer | ATGTCGCTTTCCAACAAGCTG |
|  | reverse primer | GCTCCATTGTCCAAGCAGAAT |
| Hk2 | forward primer | TGATCGCCTGCTTATTCACGG |
|  | reverse primer | AACCGCCTAGAAATCTCCAGA |
| Ldha | forward primer | TGTCTCCAGCAAAGACTACTGT |
|  | reverse primer | GACTGTACTTGACAATGTTGGGA |
| Acta2 | forward primer | GTCCCAGACATCAGGGAGTAA |
|  | reverse primer | TCGGATACTTCAGCGTCAGGA |
| Colla1 | forward primer | GCTCCTCTTAGGGGCCACT |
|  | reverse primer | CCACGTCTCACCATTGGGG |
| Timp1 | forward primer | CGAGACCACCTTATACCAGCG |
|  | reverse primer | ATGACTGGGGTGTAGGCGTA |
| SPP1 | forward primer | AGCAAGAAACTCTTCCAAGCAA |
|  | reverse primer | GTGAGATTCGTCAGATTCATCCG |
| VDR | forward primer | ACCCTGGTGACTTTGACCG |
|  | reverse primer | GGCAATCTCCATTGAAGGGG |
| CYP24A1 | forward primer | CTGCCCCATTGACAAAAGGC |
|  | reverse primer | CTCACCGTCGGTCATCAGC |
| Arg1 | forward primer | CTCCAAGCCAAAGTCCTTAGAG |
|  | reverse primer | AGGAGCTGTCATTAGGGACATC |
| Ctsl | forward primer | ATCAAACCTTTAGTGCAGAGTGG |
|  | reverse primer | CTGTATTCCCCGTTGTGTAGC |
| Slc7a2 | forward primer | TTTCCCAATGCCTCGTGTAATC |
|  | reverse primer | TGCACCCGATGACAAAGTAGC |
| GAPDH | forward primer | AGGTCGGTGTGAACGGATTTG |
|  | reverse primer | TGTAGACCATGTAGTTGAGGTCA |
| VDR promoter qPCR | forward primer | GGGAGGCGTTTACAGCAGA |
|  | reverse primer | CCTTCCGTGGAACAGTATTTGAT |
| Lyz2 Cre mice | Common | CTTGGGCTGCCAGAATTTCTC |
|  | Wild type | TTACAGTCGGCCAGGCTGAC |
|  | Mutant | CCCAGAAATGCCAGATTACG |
| Spp1-LSL-DTR-EGFP mice | Common | GGGGGCATAAGCTCTGAGAC |
|  | Wild type | CCGGAGGTGCTTACCTTCTC |
|  | Mutant | CTAGCCATGGTGCTGAGGAT |
| VDR loxp mice | Common | CTCCATCCCCATGTGTCTTT |
|  | Wild type | TTCTTCAGTGGCCAGCTCTT |
|  | Mutant | CACGAGACTAGTGAGACGTG |
